# Transcriptomic responses of the marine cyanobacterium *Prochlorococcus* to viral lysis products

**DOI:** 10.1101/394122

**Authors:** Xiaoting Fang, Yaxin Liu, Yao Zhao, Yue Chen, Riyue Liu, Qi-Long Qin, Gang Li, Yu-Zhong Zhang, Wan Chan, Wolfgang R. Hess, Qinglu Zeng

**Affiliations:** Department of Ocean Science, The Hong Kong University of Science and Technology, Clear Water Bay, Hong Kong, China; Department of Chemistry, The Hong Kong University of Science and Technology, Clear Water Bay, Hong Kong, China; Division of Life Science, The Hong Kong University of Science and Technology, Clear Water Bay, Hong Kong, China; State Key Lab of Microbial Technology, Marine Biotechnology Research Center, Shandong University, Jinan, China; Key Laboratory of Tropical Marine Bio-resources and Ecology, South China Sea Institute of Oceanology, CAS, Guangzhou, China; College of Marine Life Sciences, Ocean University of China, Qingdao, China; Genetics & Experimental Bioinformatics, Faculty of Biology, University of Freiburg, Germany; HKUST Shenzhen Research Institute, Shenzhen, China

## Abstract

Marine phytoplankton contributes to about one half of global primary production, and a significant proportion of their photosynthetically fixed organic carbon is released after viral infection as dissolved organic matter (DOM). This DOM pool is known to be consumed by heterotrophic microorganisms; however, its impact on the uninfected co-occurring phytoplankton remains largely unknown. Here, we conducted transcriptomic analyses to study the effects of viral lysis products on the unicellular cyanobacterium *Prochlorococcus*, which is the most abundant photosynthetic organism on Earth. While *Prochlorococcus* growth was not affected by viral lysis products, many tRNAs increased in abundance, which was also seen after amino acid addition, suggesting that amino acids are one of the compounds in viral lysis products that affected the expression of tRNA genes. The decreased transcript abundances of N metabolism genes also suggested that *Prochlorococcus* responded to organic N compounds, consistent with abundant amino acids in viral lysis products. The addition of viral lysis products to *Prochlorococcus* reduced the maximum photochemical efficiency of photosystem II and CO_2_ fixation while increased its respiration rate, consistent with differentially expressed genes related to photosynthesis and respiration. One of the highest positive fold-changes was observed for the 6S RNA, a non-coding RNA functioning as a global transcriptional regulator in bacteria. The high level of 6S RNA might be responsible for some of the observed transcriptional responses. Taken together, our results revealed the transcriptional regulation of *Prochlorococcus* in response to viral lysis products and suggested its metabolic potential to utilize organic N compounds.

**Importance:** Photosynthetic microorganisms called phytoplankton are abundant in the oceans and contribute to about one half of global CO_2_ fixation. Phytoplankton are frequently infected by viruses and after infection their organic carbon is released into the ocean as dissolved organic matter (DOM). Marine DOM is important for the marine food web because it supports the growth of heterotrophic microorganisms. However, the impact of viral DOM on the uninfected phytoplankton is largely unknown. In this study, we conducted transcriptomic analyses and identified many differentially expressed genes when viral DOM was added to the marine cyanobacterium *Prochlorococcus*. One effect of viral DOM is that the carbon fixation of *Prochlorococcus* was reduced by ~16%, which might affect carbon cycling in the world’s oceans since *Prochlorococcus* is the most abundant photosynthetic organism on Earth.

## Introduction

As the foundation of the ocean food web (Falkowski, 2012), phytoplankton contribute to about one half of global primary production (Field et al., 1998). A significant proportion of phytoplankton are infected by viruses (Proctor and Fuhrman, 1990; Fuhrman, 1999) and release a variety of dissolved organic matter (DOM) upon cell lysis (Kujawinski, 2011; Lønborg et al., 2013; Zhao et al., 2017; Ma et al., 2018), including carbohydrates, amino acids, and lipids (Benner and Amon, 2015). Viral lysis products have been thought to be primarily consumed by heterotrophic microorganisms (Gobler et al., 1997; Fuhrman, 1999; Azam and Malfatti, 2007; Haaber and Middelboe, 2009; Jiao et al., 2010; Sheik et al., 2014), whereas their impact on uninfected phytoplankton has historically been ignored.

The unicellular picocyanobacteria *Prochlorococcus* and *Synechococcus* are the numerically dominant phytoplankton, and they are responsible for a vast majority of primary production in many oligotrophic oceans (Liu et al., 1997; Partensky et al., 1999; Scanlan and West, 2002). The presence of viruses was reported to have a positive effect on the growth of *Synechococcus* in the ocean, suggesting that marine *Synechococcus* might benefit from the presence of viral lysis products (Weinbauer et al., 2011). Indeed, there is evidence that phytoplankton are capable of taking up organic compounds. *Prochlorococcus* and *Synechococcus* can take up amino acids (Zubkov et al., 2003; Mary et al., 2008; Gomez-Pereira et al., 2013; Bjorkman et al., 2015), dimethylsulfoniopropionate (Vila-Costa et al., 2006), and glucose (Gomez-Baena et al., 2008; Munoz-Marin Mdel et al., 2013). Natural *Prochlorococcus* populations showed rapid transcriptional responses to DOM derived from *Prochlorococcus* exudates, providing evidence that *Prochlorococcus* might use these organic compounds (Sharma et al., 2014). However, key regulators of these transcriptional responses have remained unidentified.

A widely conserved global transcriptional regulator in bacteria is the 6S RNA. In model bacteria such as *Escherichia coli* and *Bacillus subtilis*, the 6S RNA is well known to respond to changes in nutrient supply (Steuten et al., 2014; Wassarman, 2018). In *E. coli*, 6S RNA accumulates as the culture enters the stationary phase of growth and binds to the σ^70^ RNA polymerase (Wassarman and Storz, 2000). The association of 6S RNA with the σ^70^ RNA polymerase inhibits transcription at many σ^70^-dependent promoters that are used in exponential growth and activates some σ^S^-dependent promoters that are used in stationary phase (Trotochaud and Wassarman, 2004). The regulatory mechanism involves promoter mimicry, as the RNA polymerase carrying the housekeeping sigma factor σ^70^ binds to 6S RNA that mimics an open promoter complex instead of binding the respective promoter elements (Barrick et al., 2005; Cavanagh and Wassarman, 2014; Steuten et al., 2014). During recovery from the stationary phase when nutrients become available, 6S RNA serves as a template for the transcription of a ~20 nt product RNA (pRNA), leading, in turn, to its dissociation from the σ^70^ RNA polymerase (Wassarman and Saecker, 2006; Beckmann et al., 2012; Cavanagh et al., 2012). In Cyanobacteria, 6S RNAs seem to play similarly important but also divergent roles. In *Synechococcus* sp. PCC 6301, its levels change with growth, possibly due to differences in the nutrient status (Watanabe et al., 1997), although in contrast to observations in *E. coli* and *B. subtilis,* 6S RNA was abundant in exponential phase and reduced in stationary phase. Similarly, the accumulation of 6S RNA in *Prochlorococcus* MED4 was reported to be cell cycle-dependent and light-dependent (Axmann et al., 2007). Finally, genetic studies of the 6S RNA in *Synechocystis* sp. PCC 6803 and *in vivo* pull-down studies of the RNA polymerase complex demonstrated its involvement in the recovery from nitrogen depletion by accelerating the switch from alternative group 2 σ factors SigB, SigC and SigE to SigA-dependent transcription (Heilmann et al., 2017).

*Prochlorococcus* and *Synechococcus* coexist in many regions of the world’s oceans, although their distributions are not identical (Partensky et al., 1999). *Prochlorococcus* and *Synechococcus* have been shown to be actively infected by viruses (cyanophages) (Sullivan et al., 2003). In this study, we asked if the released viral lysis products could affect the uninfected neighboring cyanobacterial cells. We addressed this question by adding viral lysis products of *Synechococcus* WH8102 to *Prochlorococcus* MIT9313. These two model cyanobacterial strains were both isolated from the North Atlantic Ocean and are among the first marine cyanobacteria that have complete genomes sequenced (Palenik et al., 2003; Rocap et al., 2003). After addition of viral lysis products, we used RNA-seq to analyze the transcriptomic responses of *Prochlorococcus* MIT9313 cells, focusing on differentially expressed genes related to translation, photosynthesis, and nitrogen metabolism. We also measured whether viral lysis products affected the growth and photosynthesis of *Prochlorococcus* MIT9313. Furthermore, we found evidence that the 6S RNA may play a role in regulating the transcriptional responses of *Prochlorococcus* MIT9313 to viral lysis products.

## Results and Discussion

### *Prochlorococcus* growth is not affected by viral lysis products

We generated viral lysis products (vDOM) by infecting axenic *Synechococcus* WH8102 cultures with cyanophage S-ShM2, which does not infect *Prochlorococcus* MIT9313 (Sullivan et al., 2003). Recently, infection of *Synechococcus* WH7803 by cyanophage S-SM1 was shown to release abundant dissolved organic nitrogen compounds, including peptides derived from the major light-harvesting protein phycoerythrin (Ma et al., 2018). In our experiments, we also found that the concentrations of amino acids in vDOM were much higher than those of the natural seawater-based Pro99 growth medium (Supplementary Table 1).

To study whether vDOM affected the growth of uninfected cyanobacterial cells, vDOM was added to mid-log axenic *Prochlorococcus* MIT9313 cultures at a volume/volume ratio of 1:4, and the Pro99 growth medium was added to the control cultures (see Materials and Methods). After vDOM amendment, the cultures continued exponential growth for up to three days, with growth rates indistinguishable from the control cultures (Figure 1A). In our experiments, *Prochlorococcus* MIT9313 was grown in nutrient-replete conditions and this may explain why vDOM addition did not promote its growth. Our results were consistent with a recent study showing that the exponential growth of marine *Synechococcus* in nutrient-replete conditions was not affected after DOM was added at an amount similar to what we used here (Christie-Oleza et al., 2017).

**Figure 1.**
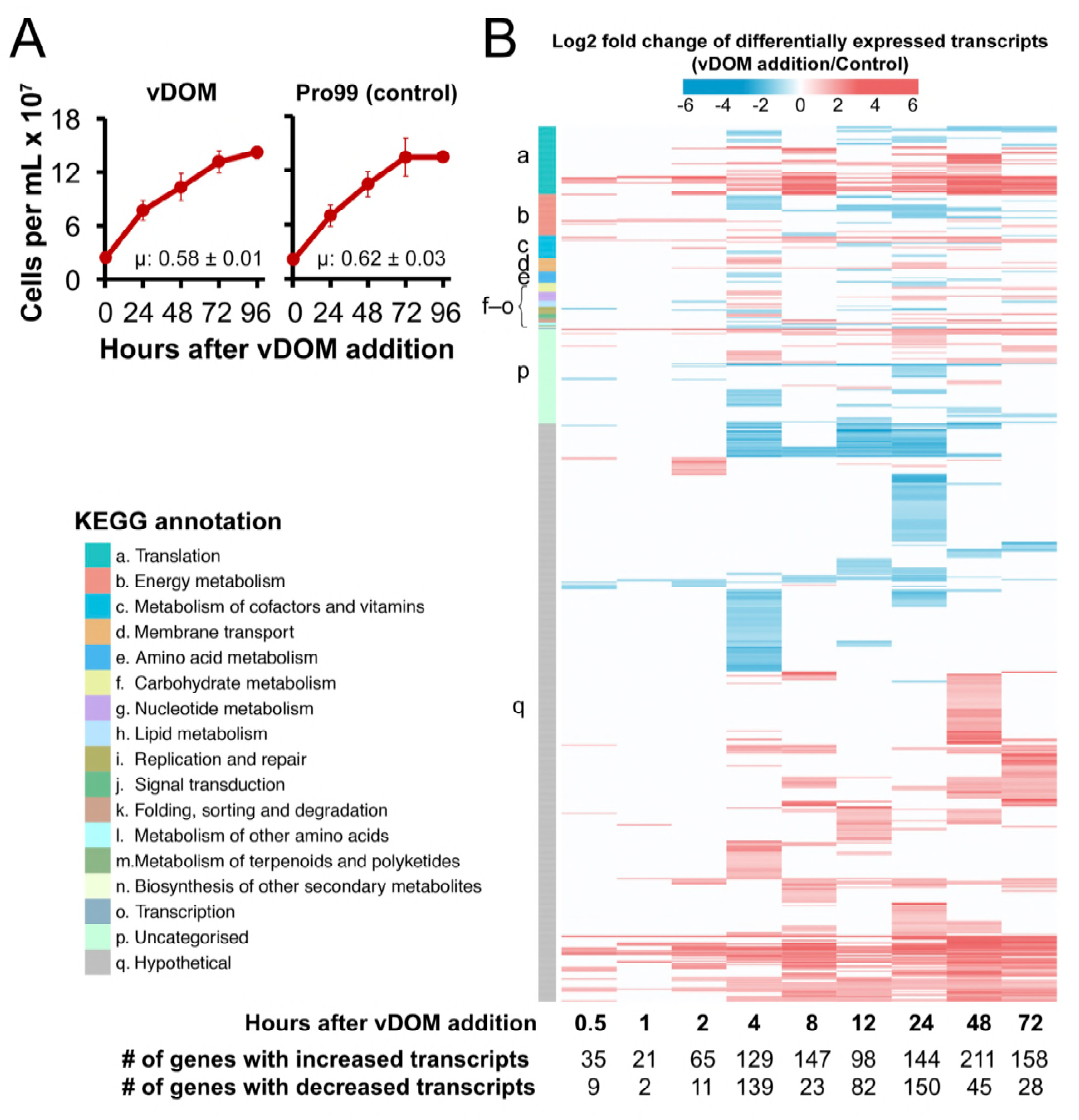
Growth and transcriptomic responses of *Prochlorococcus* after vDOM addition. (**A**) vDOM or the Pro99 growth medium (control) was added to log-phase *Prochlorococcus* MIT9313 cultures. Cell concentrations were determined by flow cytometry and were used to calculate the growth rate (µ). Data points and errors are means and standard deviations of three independent cultures, respectively. (**B**) Differentially expressed genes at 0.5, 1, 2, 4, 8, 12, 24, 48, and 72 h after vDOM addition to *Prochlorococcus* MIT9313 cells. Differentially expressed genes were identified by normalizing their transcripts in the vDOM amended cultures to those in the control cultures amended with the Pro99 growth medium (fold change ≥ 2 in either direction and an adjusted *P* value < 0.1 at least at one time point). The color bar represents log2 fold change of differentially expressed transcripts (vDOM addition/control). In the head map, a red line indicates significantly increased transcripts, a blue line indicates significantly decreased transcripts, and a white line indicates that a gene was not differentially expressed at a certain time point. Genes are grouped under KEGG (Kyoto Encyclopedia of Genes and Genomes) functional categories. The numbers of differentially expressed genes at each time point are shown below the heat map.

### An overview of the transcriptomic responses of *Prochlorococcus* to vDOM

The unchanged growth rate of *Prochlorococcus* MIT9313 after vDOM addition inspired us to wonder whether it showed any transcriptional responses. To address this question, we employed RNA-seq to analyze the transcriptomes of *Prochlorococcus* MIT9313 cells taken at 0.5, 1, 2, 4, 8, 12, 24, 48, and 72 h after amendments with vDOM or the Pro99 growth medium (control). Differentially expressed genes (Supplementary Table 2) were identified by normalizing gene expression levels in vDOM amended cultures to those of the control cultures (fold change ≥ 2 in either direction and an adjusted *P* value < 0.1 at least at one time point) (see Materials and Methods). Differentially expressed genes appeared as early as 0.5 h, indicating a rapid transcriptional response of *Prochlorococcus* cells to vDOM addition (Figure 1B).

To resolve the biological functions of these differentially expressed genes, we grouped them into functional categories (KEGG in Figure 1B and COGs in Supplementary Figure 1). Other than genes of unknown function, the most abundant functional category was translation (tRNA and ribosomal protein genes) and the second most abundant was energy metabolism (photosynthesis and respiration genes) (Figure 1B, Supplementary Figure 1). Interestingly, genes in these two categories were found to be differentially expressed when *Prochlorococcus* cells were infected by cyanophages (Lindell et al., 2007; Doron et al., 2016; Lin et al., 2016; Thompson et al., 2016). In these infection experiments, it had to remain unclear whether the transcriptional responses were caused solely by infectious phage particles, since viral lysis products were also added to the uninfected cultures. After examining genes in the functional categories membrane transport and amino acid metabolism (Figure 1B), we found that genes related to nitrogen (N) metabolism were especially affected. In the following sections, we present our detailed analysis of these differentially expressed genes.

### tRNA and ribosomal protein genes

After vDOM addition, 31 out of 43 tRNA genes showed increased transcript abundances (Supplementary Figure 2A). Not much is known about the transcriptional regulation of tRNA genes in cyanobacteria, but in *E*. *coli* cells tRNA abundances increased after amino acids were added (Dong et al., 1996). The tRNA aminoacylation level (charging level) in *E*. *coli* is also positively correlated with amino acid concentrations in the growth medium (Dittmar et al., 2005). To test whether amino acids can increase the tRNA abundances of *Prochlorococcus* MIT9313, we added glycine to *Prochlorococcus* MIT9313 and used RT-qPCR to detect the expression levels of *tRNA-Gly1*, which is the only differentially expressed tRNA gene that is long enough for qPCR measurement. Glycine did not affect the growth rate of *Prochlorococcus* MIT9313 (Supplementary Figure 3), which was previously observed in *Prochlorococcus* PCC 9511 (Rippka et al., 2000). However, similar to vDOM (Figure 2A), glycine addition increased the transcript abundances of *tRNA-Gly1* (Figure 2B), suggesting that amino acids are one of the compounds in vDOM that affected the expression of tRNA genes. Interestingly, while the tRNA abundances for 16 amino acids increased, those for aspartic acid, histidine, and tryptophan did not change significantly, and those for isoleucine decreased (Supplementary Figure 2A). This suggested that the tRNA genes of these four amino acids might be regulated by different mechanisms.

**Figure 2.**
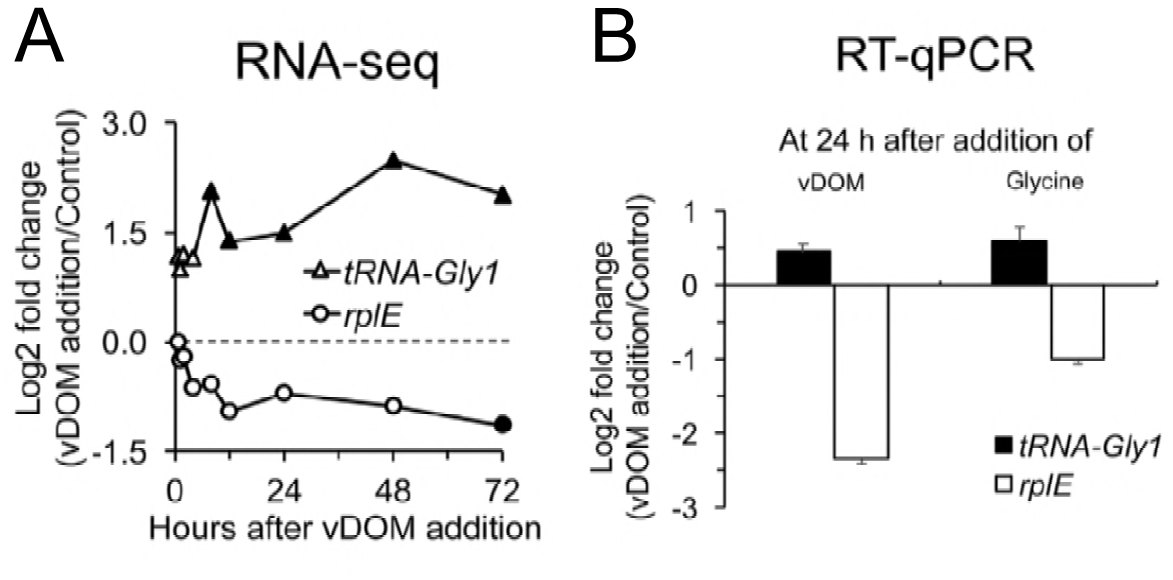
Expression of *Prochlorococcus* tRNA and ribosomal protein genes. (**A**) RNA-seq data showing the transcript abundances of the tRNA gene *tRNA-Gly1* and the ribosomal protein gene *rplE* after vDOM was added to *Prochlorococcus* MIT9313 cells. Transcript abundances in the vDOM amended cultures were normalized to those amended with the growth medium Pro99. A dotted line indicates log2 fold change = 0. Filled symbols indicate adjusted *P* values < 0.1. (**B**) RT-qPCR data showing the transcript abundances of *tRNA-Gly1* and *rplE* at 24 h after vDOM and glycine (800 µM) were added to *Prochlorococcus* MIT9313 cells. Data are mean ± s.e.m. from four biological replicates.

**Figure 3.**
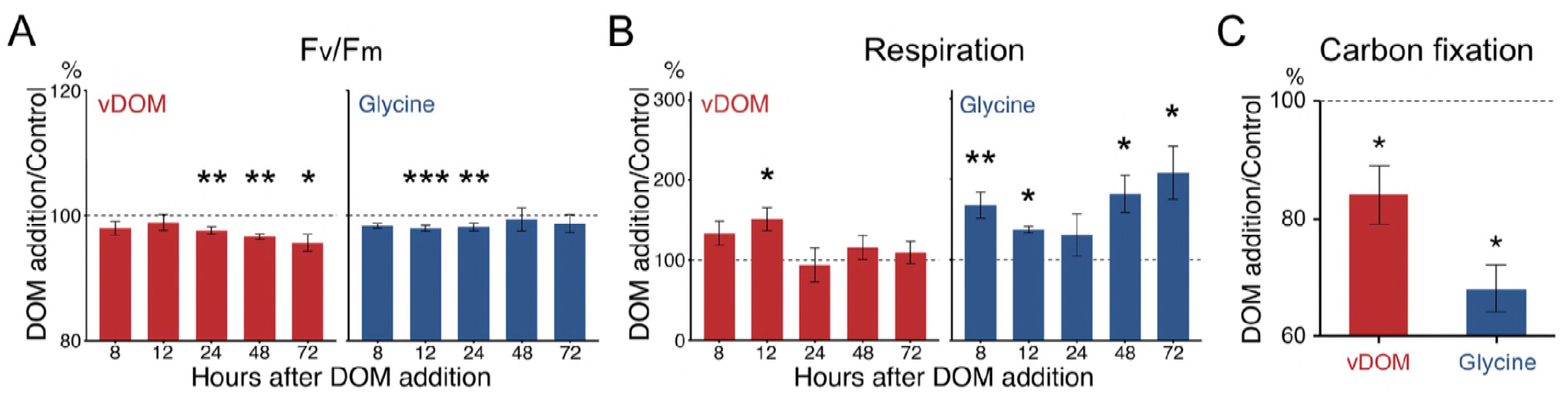
Photosynthesis and respiration of *Prochlorococcus* after DOM addition. Mid-log *Prochlorococcus* MIT9313 cultures were amended with vDOM, glycine (800 µM), or the growth medium Pro99 (control). After DOM addition, maximum photochemical efficiency of PSII (F_v_/F_m_) (**A**), respiration (**B**), and carbon fixation rate (**C**) were measured and normalized to those of the control cultures. Photosynthetic carbon fixation rates were calculated by adding NaH^13^CO_3_ into the cultures and measuring intracellular ^13^C over time (Supplementary figure 5) (Hama et al., 1983). Dotted lines indicate normalized measurement = 1. Data shown are mean ± s.e.m. from five (**A** and **B**) or three (**C**) biological replicates. Asterisks denote significant changes after DOM addition compared with the control cultures (\**P* < 0.05, \*\**P* < 0.01, \*\*\**P* < 0.001, One-way ANOVA with post-hoc Dunnett’s test).

Out of the 57 annotated ribosomal protein genes, five showed increased transcript abundances after vDOM addition, while 14 showed decreased abundances (e.g., *rplE*, Figure 2A; Supplementary Figure 2B). Amino acids might also be responsible for this transcriptional response, since glycine addition decreased the *rplE* mRNA abundance (Figure 2B). Our interpretation of the transcriptional responses of ribosomal protein genes is that they are less likely to cause a dramatic change in the number of ribosomes per cell, since vDOM or glycine addition did not affect the growth rate of *Prochlorococcus* MIT9313 (Figure 1A, Supplementary Figure 3). Recently, the relative transcript abundances of ribosomal proteins were used to assess the in situ growth rates of marine bacteria (Gifford et al., 2013), and our results suggested that this method could be affected by the DOM concentration at the sampling site.

### Photosynthesis and respiration genes

Photosynthetic and respiratory electron transport chains are important for the energy metabolism of cyanobacteria (Vermaas, 2001) and many of the respective genes were differentially expressed after vDOM addition (Supplementary Figure 4). For the light reactions of photosynthesis, transcript abundances of four photosystem II (PSII) genes and four photosystem I (PSI) genes decreased, while those of two PSII genes and three PSI genes increased (Supplementary Figures 4A, B). For genes related to chlorophyll binding/metabolism, four had increased transcripts while three decreased (Supplementary Figures 4C, D). In addition, five ATP synthase genes (Supplementary Figure 4E) and one Calvin cycle gene (Supplementary Figure 4F) showed decreased transcript abundances. For respiration, one NADPH dehydrogenase gene had decreased transcript abundances (Supplementary Figure 4G), while the succinate dehydrogenase gene *sdhB* had increased transcript abundances (Supplementary Figure 4H). We also found that three cytochrome *b*6/*f* genes had increased transcripts and three had decreased transcripts (Supplementary Figure 4I). In summary, although many photosynthesis and respiration genes were differentially expressed, their expression patterns did not clearly reveal whether the photosynthesis and the respiration of *Prochlorococcus* was affected by vDOM. Hence, we set out to measure the photosynthesis and the respiration of *Prochlorococcus* MIT9313 after vDOM addition.

**Figure 4.**
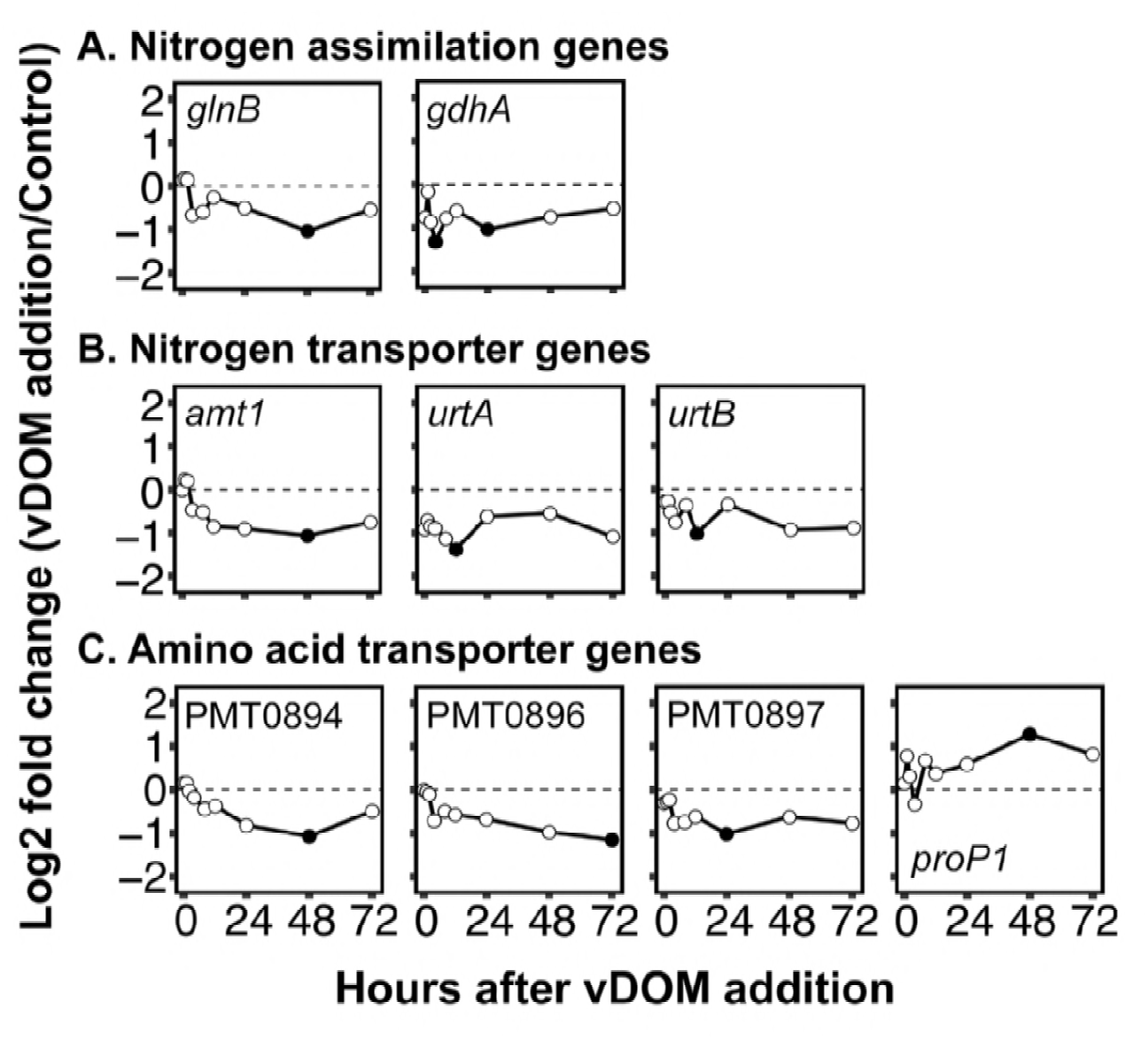
Transcript abundances of nitrogen metabolism genes. RNA-seq data showing the transcript abundances of genes for nitrogen assimilation (**A**), nitrogen transporters (**B**), and amino acid transporters (**C**). Transcripts of the DOM amended cultures are normalized to those of the control cultures amended with the growth medium Pro99. A dotted line indicates log2 fold change = 0. Filled symbols indicate adjusted *P* values < 0.1.

For photosynthesis, we measured the maximum photochemical quantum yield (F_v_/F_m_) of PSII, which reveals the potential ability of PSII to transform absorbed photons to chemical energy. F_v_/F_m_ of *Prochlorococcus* MIT9313 decreased by ~3% from 24 h after vDOM addition (Figure 3A). Despite the slight decrease of F_v_/F_m_, the carbon fixation rate of *Prochlorococcus* cells decreased by ~16% after vDOM addition (Figure 3C, Supplementary Figure 5). In contrast to photosynthesis, the respiration of *Prochlorococcus* MIT9313 cells was transiently enhanced by ~51% at 12 h after vDOM addition (Figure 3B). Similar to vDOM, glycine slightly reduced the F_v_/F_m_ of *Prochlorococcus* cells by ~2% (Figure 3A). However, glycine reduced the carbon fixation rate by ~32% (Figure 3C) and increased the respiration by up to ~108% at 72 h (Figure 3B). The more significant effects of glycine than vDOM on respiration and carbon fixation rate (Figures 3B, C) might be due to the higher DOM concentration of glycine. Although *Prochlorococcus* has been shown to take up amino acids (Zubkov et al., 2003; Mary et al., 2008; Gomez-Pereira et al., 2013; Bjorkman et al., 2015), as far as we know, our study showed for the first time that amino acids can reduce the carbon fixation of *Prochlorococcus*. With decreased carbon fixation and increased respiration, *Prochlorococcus* was supposed to accumulate less organic carbon and in that case its growth should have been inhibited. However, our results showed that *Prochlorococcus* MIT9313 maintained its growth rate after vDOM and glycine addition (Figure 1A, Supplementary Figure 3). One plausible explanation is that *Prochlorococcus* assimilates DOM compounds, which is consistent with the expression of tRNA genes (Figure 2) and is also consistent with the ability of *Prochlorococcus* to assimilate organic compounds (Zubkov et al., 2003; Vila-Costa et al., 2006; Gomez-Baena et al., 2008; Mary et al., 2008; Gomez-Pereira et al., 2013; Munoz-Marin Mdel et al., 2013; Bjorkman et al., 2015).

**Figure 5.**
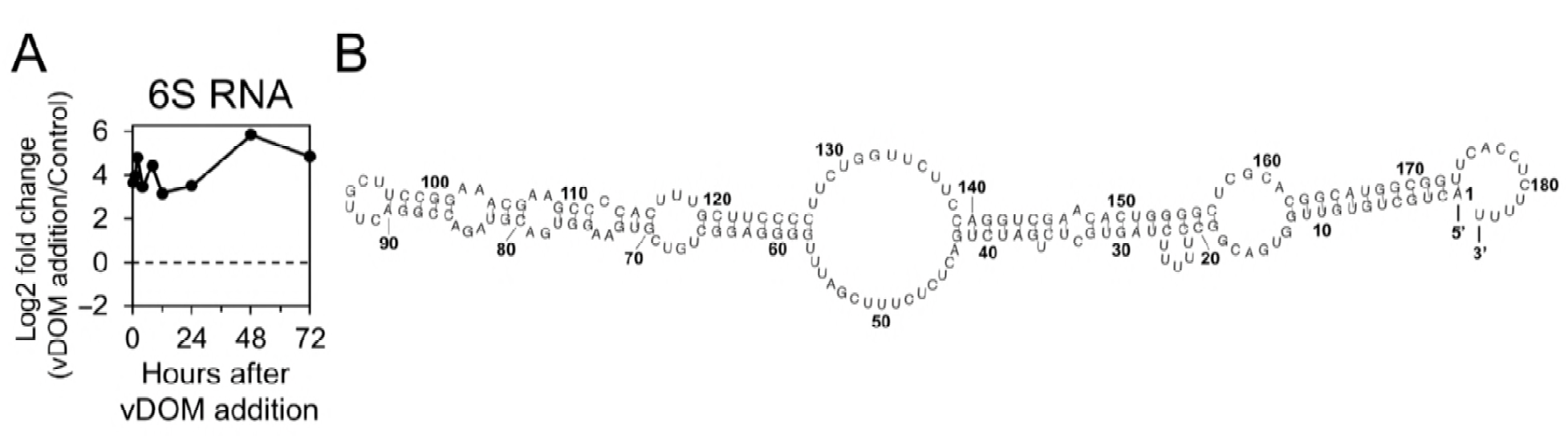
6S RNA abundance of *Prochlorococcus* MIT9313 and its predicted secondary structure. (**A**) Log2 fold change of 6S RNA (vDOM addition/control). A dotted line indicates log2 fold change = 0. All the data have adjusted *P* values < 0.1. (**B**) The secondary structure of *Prochlorococcus* MIT9313 6S RNA was predicted using the RNAfold webserver with default settings (http://rna.tbi.univie.ac.at/cgi-bin/RNAWebSuite/RNAfold.cgi) (Hofacker, 2003).

### Nitrogen metabolism genes

After vDOM addition, several genes related to N metabolism showed decreased transcript abundances (Figure 4). These genes include the global N regulator PII gene *glnB*, the glutamate dehydrogenase gene *gdhA*, and transporter genes for ammonium (*amt1*) and urea (*urtA*, *urtB*) (Figures 4A, B). As a nitrogen scavenging response, these genes show increased transcript abundances during N starvation of *Prochlorococcus* (Tolonen et al., 2006). Consistently, Sharma and colleagues observed decreased transcript abundances for ammonium and urea transporter genes when they added DOM derived from *Prochlorococcus* exudates to natural *Prochlorococcus* populations (Sharma et al., 2014). The authors of this study suggested that the DOM source they used contained labile organic nitrogen compounds that could be assimilated by *Prochlorococcus* (Sharma et al., 2014). The similar transcriptional responses of these transporter genes in our study might also be due to the organic nitrogen compounds in vDOM.

Three amino acid transporter genes (PMT0894, 0896, and 0897) also showed decreased transcript abundances after vDOM addition (Figure 4C). Although the transcriptional regulation of these genes in cyanobacteria is less clear, it has been shown in *E*. *coli* (Chubukov et al., 2014) and *Streptococcus pneumoniae* (Kloosterman and Kuipers, 2011) that bacteria down-regulate the transcription of amino acid transporter genes when amino acids are abundant in the environment (Chubukov et al., 2014). This is consistent with abundant amino acids in vDOM (Supplementary Table 1). Curiously, an amino acid transporter gene *proP* showed increased transcript abundances (Figure 4C). *E. coli* up-regulates *proP* transcription in high osmolality environments to enhance its uptake of proline and glycine betaine to avoid dehydration (Kempf and Bremer, 1998). As an adaptation to saline environments, *Prochlorococcus* MIT9313 has been found to accumulate glycine betaine and sucrose as the major compatible solutes (Klahn et al., 2010). Hence, it is possible, that the upregulated ProP system contributes to the pool of intracellularly accumulated solutes when glycine betaine becomes available in the environment, e.g. due to viral lysis of the surrounding bacteria.

### The global transcriptional regulator 6S RNA

The 6S RNA abundance increased ~13 fold at 0.5 h after vDOM addition, exhibiting the highest positive fold-change of all genes during some of the sampling points and remained highly expressed over the entire 72 h experimental period (Figure 5A, Supplementary Table 2). The 6S RNA is a non-coding small RNA (sRNA) that was first characterized in *E. coli* (Wassarman and Storz, 2000). In cyanobacteria, the 6S RNA was initially named Yfr7 and its coding gene *ssrS* is identified in all the sequenced marine cyanobacterial genomes (Axmann et al., 2005). The predicted secondary structure of *Prochlorococcus* MIT9313 6S RNA contains a central single-stranded bulge within a mostly double-stranded molecule (Figure 5B), which is conserved in cyanobacteria and *E. coli* (Rediger et al., 2012). The 6S RNA structure mimics the melted promoter DNA and it was shown in *E. coli* (Wassarman and Saecker, 2006) and *Prochlorococcus* MED4 (Rediger et al., 2012) to inhibit transcription by competing with promoter DNA binding to the RNA polymerase holoenzyme. With a conserved secondary structure (Axmann et al., 2007), the highly expressed 6S RNA of *Prochlorococcus* MIT9313 should be able to bind to RNA polymerase to regulate gene expression after vDOM addition. Interestingly, *in vivo* pull-down experiments of the RNA polymerase complex in the model cyanobacterium *Synechocystis* sp. PCC 6803 indicated that high levels of 6S RNA promote the recruitment of the housekeeping σ factor SigA, and the dissociation of alternative group 2 σ factors (Heilmann et al., 2017). *Prochlorococcus* MIT9313 harbors seven different sigma 70-type sigma factors (compared to five in *Synechocystis* sp. PCC 6803) and one type 3 alternative sigma factor (Scanlan et al., 2009). Therefore, we speculated that some of the differentially expressed *Prochlorococcus* MIT9313 genes might be attributed to sigma factor replacement mediated by the 6S RNA.

## Conclusions

Viral lysis of phytoplankton is estimated to release 6–26% of photosynthetically fixed organic carbon into the marine DOM pool (Wilhelm and Suttle, 1999). Using RNA-seq analysis, our study showed rapid and sustained transcriptomic responses of *Prochlorococcus* MIT9313 to this DOM pool. The transcriptional responses of tRNA and N metabolism genes both suggested that *Prochlorococcus* MIT9313 may take up organic nitrogen compounds in vDOM, which is consistent with previous studies that *Prochlorococcus* can take up amino acids (Zubkov et al., 2003; Mary et al., 2008; Gomez-Pereira et al., 2013; Bjorkman et al., 2015). In addition, our study showed for the first time that vDOM addition reduced the carbon fixation of *Prochlorococcus* MIT9313 by ~16%, which was also reduced by amino acid addition (~32%). This effect might be important for marine carbon cycling, considering that *Prochlorococcus* is the dominant primary producer in many oligotrophic oceans. Furthermore, our study showed that the 6S RNA was highly expressed after vDOM addition and we suggested that this global transcriptional regulator may regulate the transcriptional responses of *Prochlorococcus* MIT9313 to the availability of organic nutrients.

It should be noted that our study was conducted using the low light–adapted *Prochlorococcus* strain MIT9313 under nutrient-replete conditions. Thus, our study might only represent oceanic regions where *Prochlorococcus* cells are not severely nutrient-limited because of rapid nutrient recycling (Vaulot et al., 1995; Liu et al., 1997). Future experiments with different *Prochlorococcus* ecotypes under nutrient-limited conditions are needed, since these ecotypes have different regulatory networks (Martiny et al., 2006; Tolonen et al., 2006) and their natural distributions are shaped by various environmental factors (Johnson et al., 2006).

## Acknowledgements

This study is supported by grants to Qinglu Zeng from the National Natural Science Foundation of China (Project No. 41776132 and 41476147) and the Research Grants Council of the Hong Kong Special Administrative Region, China (Project No. 16102317 and 16103414). This study is partially supported by grants to Yu-Zhong Zhang from the National Natural Science Foundation of China (Project No. 31630012 and U1706207). We thank Dr. Yehui Tan and Jiaxing Liu in the South China Sea Institute of Oceanology and Dr. Dinghui Zou in the South China University of Technology for their experimental assistance.

## Author Contributions

Q.Z. and X.F. designed the project; X.F. performed all of the experiments and data analysis, with assistance from other authors; Y.L. measured DOC, DN, *tRNA* and *rplE* expression; Y.Z. and W.C. did amino acid measurements; Y.C. measured *tRNA* and *rplE* expression; R.L. measured growth rates and did axenicity tests; G.L. measured F_v_/F_m_ and respiration; Q.-L.Q. and Y.-Z.Z. analyzed genes related to amino acid uptake; Q.Z. and X.F. wrote the manuscript with contributions from all authors.

## Legend for Supplementary Materials

**Supplementary Figure 1. Differentially expressed *Prochlorococcus* MIT9313 genes categorized in clusters of orthologous groups (COGs)**

Differentially expressed genes were identified at 0.5, 1, 2, 4, 8, 12, 24, 48, and 72 h after vDOM addition to *Prochlorococcus* MIT9313 cells. In the head map, a red line indicates increased transcripts, a blue line indicates decreased transcripts, and a white line indicates that a gene was not differentially expressed at a certain time point. Genes are grouped bases on their COGs categories.

**Supplementary Figure 2. Transcript abundances of tRNA (A) and ribosomal protein genes (B)**

Heat map shows the relative transcript abundances of the vDOM amended cultures normalized to those of the control cultures amended with the growth medium Pro99. The color bar represents log2 fold change of transcripts (vDOM addition/control). Genes with increased, decreased, and not differentially abundant transcripts are shown by red, blue, and grey boxes, respectively. The tRNA genes in **A** are grouped based on the side chains of their corresponding amino acids.

**Supplementary Figure 3. *Prochlorococcus* MIT9313 growth after glycine addition**

The growth medium Pro99 (control) or glycine (800 µM) were added to N-replete log-phase *Prochlorococcus* MIT9313 cultures. Cell concentrations were measured by flow cytometry and were used to calculate the growth rates (µ). Error bars indicate the standard deviations of three biological replicates and are smaller than the data point when not apparent. Note that *Prochlorococcus* MIT9313 showed different growth rates in the Pro99 media used in this figure and in Figure 1A, which were made of natural seawater collected in different seasons.

**Supplementary Figure 4. Transcript abundances of photosynthesis and respiration genes**

RNA-seq data shows log2 fold change of transcripts (vDOM addition/control). A dotted line indicates log2 fold change = 0. Filled symbols indicate adjusted *P* values < 0.1.

**Supplementary Figure 5. Carbon fixation rates of *Prochlorococcus* after DOM amendments**

Mid-log *Prochlorococcus* MIT9313 cultures were amended with vDOM, glycine or the Pro99 culture medium. At the same time, NaH^13^CO_3_ was also added to the cultures. Photosynthetic production (fg C cell^−1^) was calculated using the ^13^C method (see Materials and Methods). Since photosynthetic production increased linearly over incubation time (*r*^2^ > 0.9 in all the cultures), photosynthetic carbon fixation rates (fg C cell^−1^ h^−1^) were estimated by linear-fitting photosynthetic production against incubation time and were shown in each graph. Data shown are from three biological replicates.

**Supplementary Table 1. Concentrations of total combined amino acids**

The total concentrations of several amino acids are shown in groups because of overlapping signals during HPLC-FLD measurements (leucine, isoleucine, and phenylalanine in one group, and glutamine and asparagine in another group). Concentrations of alanine, cysteine, proline, and tyrosine could not be measured using our method due to the low resolutions of individual amino acid standards. Data are mean ± standard deviation of four biological replicates. In the Pro99 growth medium, amino acids were not detected (ND).

**Supplementary Table 2. Differentially expressed *Prochlorococcus* MIT9313 genes**

Transcript abundances of the vDOM amended cultures were normalized to those of the control cultures amended with the Pro99 growth medium. For each log2 fold change, * indicates a significant change (adjusted p-value < 10%, and log2 fold change bigger than 1 or smaller than - 1).

**Supplementary Table 3. RNA-seq libraries constructed in this study**

**Supplementary Table 4. Number of mapped reads per gene**

**Supplementary Table 5. qPCR primers used in this study**

## Materials and Methods

### Cultivation of *Prochlorococcus* and *Synechococcus*

Axenic *Prochlorococcus* MIT9313 and *Synechococcus* WH8102 cultures were grown in polycarbonate bottles (Nalgene) with the Pro99 medium (Moore et al., 2002), which was based on natural seawater from Port Shelter, Hong Kong. *Prochlorococcus* and *Synechococcus* cultures were incubated at 24°C under continuous cool white light at 15–20 and 30 μmol quanta m^−2^ s^−1^, respectively. Bulk culture chlorophyll fluorescence was monitored using a fluorometer (10-AU model, Turner Designs). For cell counting, 200 μL culture was preserved with 1% Glutaraldehyde solution (50 wt. % in H_2_O, SIGMA) and stored at –80°C until use. Preserved cells were counted using a flow cytometer (BD FACSCalibur) with the ModfitLT software.

### Axenicity tests

*Prochlorococcus* and *Synechococcus* cultures were routinely tested for axenicity (no contamination of heterotrophs) by inoculating them in three purity broths ProAC (Morris et al., 2008), MPTB (Saito et al., 2002), and ProMM (Berube et al., 2015). The cyanobacterial cultures were considered axenic when no purity broth became turbid within one week at room temperature. Axenicity was also tested by flow cytometry and epifluorescence microscopy. In axenic cultures, all of the SYBR-Gold staining cells were *Prochlorococcus* or *Synechococcus* (determined by their autofluorescence), and no other SYBR-Gold staining cells were observed.

### Preparation of viral lysis products (vDOM)

vDOM was generated by infecting early-log *Synechococcus* WH8102 cultures (~3 × 10^7^ cells per mL) with 1/10 cyanophage S-ShM2 lysate (volume/volume) until cultures became clear. Lysates were filtered through a 0.2 µm filter to remove cell debris and then were stored in acid-washed glassware at 4°C in the dark for several days before they were used in vDOM addition experiments.

### Measurements of dissolved organic carbon and dissolved nitrogen

The dissolved organic carbon and dissolved total nitrogen (inorganic and organic) concentrations of the DOM sources were measured and analysed using an automated Shimadzu TOC analyser (TOC-V_*CPH*_ and Autosampler ASI-V, Shimadzu, Japan) according to the manufacturer’s instruction. Dissolved organic carbon concentrations in vDOM and Pro99 were 682.95 ± 14.88 and 367.22 ± 5.24 μM, respectively. Dissolved total nitrogen concentrations in vDOM and Pro99 were 804.59 ± 14.64 and 870.54 ± 12.18 μM, respectively.

### Measurements of total combined amino acids

Dissolved proteins in vDOM were concentrated using a centrifugal filter unit (10 KDa cutoff, Amicon). The resulting protein concentrates from 50 mL vDOM were dissolved in ~300 µL ultrapure water, and were hydrolyzed using 5 μg protease XIV (Sigma) at 37°C for 16 h as described previously (Liu et al., 2016). Protein hydrolysates were then derivatized with 9-fluorenylmethyl chloroformate (Fmoc-Cl) and individual amino acid concentrations were determined by high performance liquid chromatography coupled with fluorescence detection (HPLC-FLD) following published protocols (Buha et al., 2011; Liu et al., 2016). The concentrations of amino acids in the Pro99 growth medium were measured using the same method and were all below the detection limit (Supplementary Table 1).

### vDOM addition into *Prochlorococcus* MIT9313 cultures

In the oceans, ~15% of cyanobacteria are infected by cyanophages at any given time (Proctor and Fuhrman, 1990), and hence ~85% uninfected cyanobacteria are exposed to the DOM released by the infected cells. To mimic this effect, vDOM was added to mid-log phase *Prochlorococcus* MIT9313 cultures at a volume/volume ratio of 1:4 and the cultures were then incubated under continuous light. In the control cultures, the growth medium Pro99 was added at a 1:4 ratio.

### RNA-seq library construction

With two biological replicates for each DOM amendment, cultures were collected at 0.5, 1, 2, 4, 8, 12, 24, 48, and 72 h after DOM addition. About 40–80 mL *Prochlorococcus* culture was spun down at 15,000 g for 15 min at 4°C, and cell pellets were flash frozen in liquid nitrogen and stored at –80°C. The mirVana RNA isolation kit (Ambion) was used to extract total RNA from cell pellets and the Turbo DNA-free kit (Ambion) was used to remove residual genomic DNA. Total RNA was concentrated with the RNA Clean & Concentrator-5 kit (Zymo Research), 150 ng total RNA was then fragmented into 60–200 nt by magnesium catalysed hydrolysis (40 mM Tris-Acetate, pH 8.1, 100 mM KOAc, and 30 mM MgOAc) for 4 min at 83°C, and purified with the RNA Clean & Concentrator-5 kit. As we described previously (Lin et al., 2016), strand-specific RNA-seq libraries were constructed with a dUTP second-strand marking protocol, while16S and 23S cDNA molecules were degraded by a duplex-specific nuclease (DSN) treatment. After DSN treatment, Illumina sequencing primers with barcodes were used to amplify the libraries. Equal amounts of 36 barcoded libraries were pooled within one lane for a total of four lanes, and paired-end sequencing was done by Illumina HiSeq 2000 (49 nt for insert + 6 nt for barcode).

### Data availability

The raw reads of RNA-seq data have been submitted to the European Nucleotide Archive (http://www.ebi.ac.uk/ena) under project number PRJEB22768 (ERP104472). Sample accession numbers are listed in Supplementary Table 3.

### RNA-seq data analysis

After Illumina sequencing, reads were separated based on their barcodes and their quality was assessed by FastQC (www.bioinformatics.babraham.ac.uk/projects/fastqc). The resulting clean reads were mapped against the *Prochlorococcus* MIT9313 genome using the Burrows-Wheeler Aligner (BWA) (Li and Durbin, 2009). HTSeq (Anders et al., 2015) was then used to calculate the number of reads perfectly aligning to the sense and antisense strands of ORFs, rRNAs, tRNAs, and intergenic regions. For reads spanning two ORFs, they were counted once for each ORF. Reads that mapped to the 16S and 23S rRNA genes were removed manually from the total reads before further analysis. Among the 2,765 annotated genes, over 98.8% were found to be transcribed in at least one sample, and 93.2% of the transcribed genes had a sequencing depth of more than 10 times, which indicated a thorough coverage. For each sample, the numbers of mapped reads per gene are listed in Supplementary Table 4. The two biological replicates at each time point were highly reproducible (Pearson’s R value > 0.95 at most time points, Supplementary Table 3).

### Identification and functional categorization of differentially expressed genes

The DESeq2 package (Love et al., 2014) in R (www.R-project.org) was used to identify differentially expressed genes with default parameters. For each sample, DESeq2 first normalized the number of reads per gene to the total number of mapped non-rRNA reads in that sample. DESeq2 then compared the normalized gene expression levels in the DOM amended samples to those in the control samples amended with the growth medium Pro99. We considered a gene as differentially expressed after DOM amendment if its transcript abundances showed a fold change ≥ 2 in either direction and an adjusted *P* value < 0.1. Adjusted *P* values were calculated by DESeq2 using the Benjamini-Hochberg procedure (Love et al., 2014). Differentially expressed genes are listed in Supplementary Table 2.

The Kyoto Encyclopedia of Genes and Genomes (KEGG) database of *Prochlorococcus* MIT9313 was fetched on-line (www.kegg.jp/kegg-bin/get_htext?htext=pmt00001) (Kanehisa et al., 2017). The Clusters of Orthologous Groups (COGs) were acquired by mapping *Prochlorococcus* MIT9313 protein sequences against the NCBI COG database (www.ncbi.nlm.nih.gov/COG/) (Tatusov et al., 1997). In total, 1,108 COG-assigned proteins were found, with an E-value cutoff of 0.01. Higher categories of KEGG and COG classes were used to cluster the differentially expressed genes of each DOM amendment using the pheatmap package in R (Kolde, 2015) (CRAN.R-project.org/package= pheatmap).

### Quantitative reverse transcription PCR

In Figure 2B, vDOM, glycine (diluted in Pro99), or Pro99 was added to mid-log *Prochlorococcus* MIT9313 cells at a volume/volume ratio of 1:4. The final glycine concentration was 800 µM. At 24 h after DOM addition, cells were collected by centrifugation. The relative transcript abundances of *tRNA-Gly1* and *rplE* genes were measured by quantitative reverse transcription PCR (RT-qPCR) following a previous protocol (Lin et al., 2016). Briefly, after RNA extraction and reverse transcription, cDNA copies were quantified using a QuantiTect SYBR Green PCR Kit (QIAGEN) with 0.5 μM forward and reverse primers (see Supplementary Table 5 for primer sequences). The relative transcript abundances of *tRNA-Gly1* and *rplE* genes were normalized to those of the host *rnpB* gene (Lindell et al., 2007; Zeng and Chisholm, 2012).

### Carbon fixation rate

vDOM, glycine (diluted in Pro99), or Pro99 was added to mid-log *Prochlorococcus* MIT9313 cells at a volume/volume ratio of 1:4 (this was also done before the measurements of F_v_/F_m_ and respiration). The final glycine concentration was 800 µM. Immediately after DOM amendments, freshly prepared 1 M NaH^13^CO_3_ was added to the cultures at a final concentration of 6 mM. At 0, 4, 8, 12 and 24 h after NaH^13^CO_3_ addition, 50 mL culture was centrifuged at 4°C at 20,000 g for 5 min. Cell pellets were washed twice with 50 mL Milli-Q water to remove extracellular NaH^13^CO_3_. Cell pellets were then transferred to a 1.5 mL Eppendorf tube and subjected to freeze-dry for 1–2 h. The freeze-dried pellet was washed with 200 μL hydrochloric acid (5%) for less than 5 min to remove intracellular inorganic carbon. Acid was removed after centrifugation and the cell pellet was washed twice by ultrapure water (200 μL water for the first wash and 400 μL for the second). Cell pellets were freeze-dried, weighted, and wrapped in tin capsules. ^13^C and total carbon measurements were done by the Stable Isotope Facility of the University of California at Davis.

Photosynthetic carbon fixation rate was calculated based on the intracellular ^13^C measurements (Hama et al., 1983). Briefly, the photosynthetic carbon fixation rate *P* (fg C cell^−1^ h^−1^) can be obtained using the equation below:

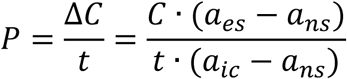

Here, *a*_*es*_, *a*_*ns*_ and *a*_*ic*_ are the atom percentages of ^13^C over total carbon in the experimental samples, natural organic samples, and natural total inorganic carbon, respectively. *C* is the particulate organic carbon (POC) in the experimental samples (fg C cell^−1^), *ΔC* is POC increase during NaH^13^CO_3_ incubation (fg C cell^−1^), and *t* is the incubation time in hours. After addition of vDOM, glycine, or Pro99, intracellular organic carbon of *Prochlorococcus* MIT9313 increased linearly throughout the experimental period (Supplementary Figure 5). Carbon fixation rates were estimated by linear-fitting the calculated photosynthetic production against incubation time (Supplementary Figure 5). The estimated carbon fixation rates in our study (0.52–0.62 fg C cell^−1^ h^−1^) are comparable with those of *Prochlorococcus* lab cultures and field populations using similar methods (Partensky et al., 1999).

### Photosystem II photochemical efficiency

The maximum PSII photochemical efficiency of mid-log *Prochlorococcus* MIT9313 was measured using a fluorometer (PSI FL 3500, Photon Systems Instruments, Czech Republic) with the fast repetition rate (FRR) fluorescence technique (Kolber et al., 1998). Prior to each measurement, 2 mL culture was loaded into a 10 × 10 mm cuvette and dark-adapted for 10 min, and the FRR induction was driven by a train of 40 × 1.2 µs flashlets (625 nm, ~100,000 µmol photons m^−2^ s^−1^). The resulting FRR induction curves were then analysed in R with a published model (Kolber et al., 1998) to derive F_0_, the base line fluorescence of cells after 10 min darkness, and F_m_, the maximal fluorescence with all PSII closed, respectively (van Kooten and Snel, 1990). The maximum photochemical quantum yields of PSII (F_v_/F_m_) was then calculated by (F_m_ − F_0_) / F_m_. All measurements were conducted at room temperature under the dark.

### Measurement of respiration rate

The respiration rate was obtained by measuring the oxygen consumption of *Prochlorococcus* MIT9313 in the dark, using a FireSting Optode sensor controlled through the Oxygen Logger software (PyroScience, Germany). Prior to measurements, 10 mL culture was loaded into a chamber in a customized acrylic vial, which is connected to a circulating thermostatted bath (Cole-Parmer, U.S.A.) to maintain the growth temperature. A magnetic stir bar was put into the chamber to remove air bubbles. The oxygen concentration in the cultures was continuously monitored (1-s interval) in the dark for 5 min at growth temperature. Oxygen removal rates were calculated in R by linear-fitting oxygen concentration (μmol O_2_ L^−1^) against elapsed time (s). Respiration rates (fg O_2_ cell^−1^ h^−1^) were then calculated by normalizing the oxygen removal rate to total number of cells (cells per 10 mL).

